# Universal phylogenetic inertia in body temperature evolution across endothermic and ectothermic tetrapods

**DOI:** 10.1101/2025.10.27.684885

**Authors:** Eliza Tarimo, Emma White, Brooke L. Bodensteiner, Martha M. Muñoz, Brian P Waldron, Josef C. Uyeda

## Abstract

Species must adapt to persist in a changing world. As global temperatures rise, how species adapt and respond to thermal shifts is crucial for anticipating global patterns of biodiversity change. Land vertebrates can be divided into two major thermoregulatory strategies, endothermy and ectothermy. One might hypothesize that, given their reputation as being “cold blooded,” ectotherms are thermal generalists, capable of operating across a greater range of body temperatures than endotherms and exhibit greater plasticity and evolvability in body temperature. However, a wide variety of traits and ecologies could modulate responses of thermal physiology to environmental change. Here, we employ macroevolutionary models to estimate the rate of adaptation of thermal physiology across squamates, mammals, and birds in the context of their ecology, physiology, and changing climatic conditions and whether there are fundamental differences in how the three clades respond to their environments. We find stronger relationships between squamates’ body temperature and their environment than in birds and mammals, significant effects of diel activity (nocturnal and diurnal) on body temperature evolution in all clades, and no effect of aquatic/terrestrial habits and rumination on the evolution of body temperature in mammals. Most surprisingly, our findings suggest shared limits on the evolution of thermal physiology across ectothermic and endothermic groups that argue for universal constraints on the rate of evolution in thermal physiology while explaining disparate patterns of body temperature and niche evolution across groups.

## Introduction

The water-to-land transition was a major milestone in the evolution of tetrapods, enabling access to terrestrial habitats and giving rise to tens of thousands of modern species. Over the past 350+ million years of terrestrial evolution, tetrapods have diversified from pole to pole, expanding to all continents, and from sea level to several kilometers above it. This vast geographic expansion has exposed them to a remarkable range of thermal conditions, with environmental temperatures spanning nearly 100°C (−67.7 to 50°C; Clarke, 2014). Nevertheless, body temperature (or preferred/active body temperature in ectotherms) of the major tetrapod clades generally lies between the much narrower range of 20 to 45 °C.

George Williams (1992) long puzzled over the surprisingly limited range of tetrapod body temperatures, and in particular the even smaller range observed in endotherms. Given the potential benefits of thermal adaptation to prevailing climate and the remarkable diversity in other phenotypic traits that accumulate in the same clades, the apparent evolutionary stasis in body temperature constitutes a paradox (Williams, 1992). Why should mammalian adaptive radiation involve rapid divergence in body size and shape, for example, but not in core temperature? Probing the causes of thermal conservatism may reveal fundamental asymmetries in the evolvability of phenotypic diversity. Moreover, better understanding the phylogenetic pattern and ecological predictors of body temperature conservatism is of key importance for predicting impacts of global change, as lineage-specific constraints on adaptive responses to changing temperatures could leave some species more vulnerable than others to rising temperatures.

Evolutionary stasis can arise from a variety of (non-mutually exclusive) limitations on adaptation, including stabilizing selection, behavioral buffering, and genetic constraints, but seem especially likely to emerge from underlying biochemical constraints (Tattersall et al., 2012, Somero et al. 2017). The upper thermal limit of enzyme function is tightly constrained at high temperatures, although relatively fewer constraints establish the lower boundary of biochemical performance at cold temperatures (Tattersall et al., 2012, Somero et al. 2017, Angilletta et al., 2010). Understanding the differences in temperature adaptation between groups of species over macroevolutionary time, and how these responses are shaped by their unique and shared adaptive strategies, could provide a better understanding of how different species respond to environmental change. Previous phylogenetic studies have found distinct evolutionary dynamics for body temperature between endotherms and ectotherms. Among terrestrial vertebrates, for example, Moreira et al. (2021) found that the rate of body temperature evolution was twice as fast for ectotherms than for endotherms. Qu & Wiens (2020) found a weak correlation between climatic niche variables and physiological parameters such as body temperature in squamates, birds, and mammals (all R^2^<0.03), with the strongest correlation in amphibians (R^2^=0.25). These differences highlight that evolutionary responses to the thermal environment varies among clades, and broader comparisons may provide insight into the sources of stasis in body temperature evolution.

At the macroevolutionary scale, evolutionary conservatism in traits is often assessed by examining a trait’s phylogenetic signal, or the tendency for closely related species to resemble each other more than to distant relatives (Pagel, 1999). However, making evolutionary inferences from phylogenetic signal is not always straightforward (Revell et al. 2008). Phylogenetic signal in traits such as body temperature can arise from shared responses to similar environments/ habitat that themselves may change according to processes that generate phylogenetic signal (Freckleton & Jetz, 2009; Hansen & Orzack, 2005) or from a lag in adaptation, sometimes termed ‘phylogenetic inertia’ (Hansen et al. 2008). Phylogenetic inertia occurs when current trait values have not reached their long-term adaptive optima under a given environment. The presence of adaptation does not negate the presence of inertia, nor does inertia imply the absence of adaptation; both contribute to observed evolutionary patterns, necessitating a model that assesses them jointly while controlling for other variables. Although many studies have examined body temperature evolution and its relationship to environmental factors, fewer have attempted to separate phylogenetic inertia, or the ‘slowness’ of adaptation, from the effects of differential rates of change in the climatic variables species experience (Labra et al., 2009; Hansen & Orzack, 2005; Hansen et al., 2008; Hansen & Bartoszek, 2012; Bodensteiner et al. 2021).

Besides the underlying evolvability of body temperature itself, the behavior of organisms may also contribute to phylogenetic lags in thermal adaptation. Organisms can use behavioral thermoregulation to maintain their body temperature within a certain range and limit exposure to thermal extremes; across environmental gradients, such buffering behavior can limit local adaptation and slow the rate of physiological evolution, a phenomenon termed the ‘Bogert effect’ (Bogert 1949; Huey et al. 2003; Muñoz 2022). Thermoregulation, however, is not an equally accessible strategy for all organisms; access to greater thermal variation affords more opportunities for behavioral buffering. For example, there is greater thermal variation during the day than at night (Ghalambor et al. 2006; Tattersall et al., 2012; Muñoz and Bodensteiner 2019), so the Bogert effect is commonly observed in diurnal than in nocturnal organisms (Muñoz, 2022). Likewise, airborne habitats are generally more thermally variable than aquatic habitats (i.e., higher specific heat capacity in water), so the Bogert effect should be more readily observed in terrestrial than in aquatic species (Bodensteiner et al. 2021). While the Bogert effect is typically viewed as modulating the evolutionary rate of physiological traits, it is also possible that fundamental constraints on the evolvability of physiology necessitate these alternative modes of adaptation— essentially, that the alternative to behavioral adaptation is not physiological adaptation, but rather a failure to adapt entirely if physiological adaptation is highly constrained. Because endotherms metabolically regulate their body temperature without much influence from their environments, an informative test is to compare the phylogenetic inertia between endotherms and ectotherms, as well as its responsiveness to predictors of behavioral thermoregulation, such as diel activity.

Here, we ask how species adapt to environmental temperatures by using models that quantify how body temperature responds to a set of adaptive factors hypothesized to be strong influences on thermal physiology, including behavioral and diel patterns, environmental variables, and evolutionary history (Labra et al. 2009). Building on previous studies (Qu & Wiens 2020, Moreira et al. 2021), we examine this question broadly across major vertebrate groups that differ in their mode of thermoregulation, either endothermy or ectothermy. However, unlike previous studies we examine these factors in the adaptation-inertia framework to jointly account for and estimate phylogenetic inertia in body temperature, which we hypothesize may be significant in thermal physiology. Testing these macroevolutionary relationships in a framework that accounts for adaptive evolution (Hansen et al. 2008), and how these dynamics differ between clades, is of critical importance for making sense of how and why species adapt--or fail to adapt--to changing environments.

## Methods

All analyses were done in R. We incorporated phylogeny, environmental variables, and organismal traits in a phylogenetic regression framework to examine their effect on T_b_. We estimated adaptation and inertia using Hansen et al.’s (2008) adaptation-inertia framework, *slouch*, which estimates the regression between environments and body temperature while assessing inertia through the phylogenetic signal in residual variation. Using model selection, we identified which predictors jointly best explain patterns of T_b_ evolution.

To estimate the lag in adaptation, *slouch* provides an estimate of the “optimal” regression, which is the relationship between variables that would exist given an infinite amount of time for adaptation in a stable environment; in other words, the optimal regression describes a pattern of optimal adaptation of body temperature to environment. We then compared the evolutionary (*i.e*., observed) regression to the optimal regression. If there is no phylogenetic inertia, then the evolutionary and optimal regressions will converge to the same relationship. By contrast, a significant phylogenetic lag in adaptation will result in differences in slope between the optimal and evolutionary regressions. We measure inertia as phylogenetic half-life, which is the time it takes for the evolutionary traits to evolve halfway to their optimal states in millions of years (my). This value, calculated simultaneously by the model, is inversely related to the rate of adaptation (*ln 2/* α).

Climatic variables interact extensively to shape the realized climate experienced by organisms, which in turn can influence their physiology and thermoregulation (Rozen‐Rechels et al., 2019). Therefore, besides the direct effects of environmental temperature, we include precipitation as a predictor. For example, water availability (or lack thereof) could alleviate the costs of very warm temperatures on the fitness of an organism. We used mean annual temperature (*T*_*a*_) and mean annual precipitation (*P*_*mm*_) as our primary environmental predictors to understand how body temperatures track and adapt to changing environmental temperature after accounting for other significantly covarying factors. To test the hypothesis that behavioral thermoregulation, we also include diel activity (nocturnal and diurnal). Specifically, our prediction is that the evolutionary lag (*i.e*., phylogenetic half-life) imposed by behavior will be stronger in diurnal than nocturnal species (Muñoz and Bodensteiner 2019). In our analysis, we aimed to see if these differences in diel activity influence T_b_ optima by comparing rates of T_b_ evolution between endotherms and ectotherms.

Additionally, we account for other variables such as aquatic vs terrestrial habitat (Rozen‐Rechels et al., 2019), which strongly effects the thermal environment and impacts organisms’ thermal sensitivity, physiology and behavior (Clusella-Trullas et al., 2011). Preliminary exploration of our compiled datasets suggests that few aquatic birds or squamates are present in the data to test this hypothesis in those clades, so we limit our inclusion of habitat predictors to mammals. Additionally, within mammalian ruminants, *i.e* members of the suborder Ruminantia, excluding the Tragulidae family, experience elevated heat-generation from microbial metabolic activities associated with rumination and exhibit counter-adaptations in response. For example, a unique structural cooling adaptation known as the carotid rete mirabile forms an intricate network of blood vessels that plays a vital role in regulating brain temperature from that of the core (Daniel & Dawes, 1953; Ferreira et al., 2021; Fuller et al., 2016). Given the evidence of selection and adaptation in the thermal biology of this group, we hypothesize that there is significant potential for selection and adaptation in ruminant body temperature as well. To test this hypothesis, we include a clade-level factor for Ruminantia to allow for unique effects on body temperature relative to that of other mammalian clades.

### Thermal physiology data

We assembled a global dataset of squamates, mammals, and birds. Body temperature data from 857 mammal and 476 bird species were downloaded from physiological reference values from the Species360 database (Species360, 2014), which is standardized by health status and measurement method. We acquired field body temperature data for 501 squamate species from (Bodensteiner et al., unpublished) where each species had body temperature measured from behaviorally active lizards, along with standard deviations where possible. We only included field measured body temperature that followed the following criteria: 1) thermal imaging camera or IR gun data was included if the lizard was <3 grams or the authors provided some correction to their data to account for thermal inertia associated with larger body sizes. 2) Data was excluded if the lizards were shot with a bb gun before body temperature was measured. 3) Data was excluded if it took longer than 10 seconds to measure body temperature from time of capture. We associate these temperatures to the T_b_ values in endotherms that maintain body temperature values through physiological means via the assumption that actively thermoregulating lizards will be near their preferred values, which we view as conceptually equivalent as the average body temperature of endotherms. All datasets also include at least partial data on standard error values for most measurements. of body temperature (i.e. we accounted for the higher uncertainty in body temperature in ectothermic squamates than in endotherms). We replaced all missing standard errors with the mean standard error value for body temperature in each dataset to account for any missing standard error values, and square all standard errors to obtain measurement variances.

### Diel activity and habitat use data

We used diel activity state (diurnal vs. nocturnal) data for squamates, mammals and birds from a compilation by Moreira et al. (2021). While activity patterns likely follow continuous underlying variation in environmental temperatures, we analyzed these as binary nocturnal and diurnal states as a practical matter, and to maximize sample sizes within groups. Species that fell in crepuscular (n=6) and arrhythmic (n=20) species were coded in nocturnal and diurnal states, respectively. Aquatic vs. terrestrial mammalian states were gathered from Brooks et al. (2024), which was compiled from Myhrvold et al. (2015).

### Rumination

Our dataset contains body temperatures for 89 species of ruminants from all the six families they span. We include a binary predictor for rumination in this monophyletic group and analyzed and compared body temperature evolution of this clade with the rest of the mammalian species. While rumination is a singular event that can be confounded by other shared features of ruminants (Uyeda et al. 2018), our primary goal was to allow for a ruminant effect when examining patterns of thermal adaptation broadly across mammals.

### Ecological/Climatic Data Collection

All species occurrences for squamates and mammals were requested and downloaded from the Global Biodiversity Information Facility (GBIF, 2023). Species occurrences were then imported into the programming software, R, using the function “occ_download_get” from the package *rgbif* (Chamberlain et al. 2021). After all species occurrences were imported, we subsetted for species’ scientific names, longitudes, latitudes, and each species country code. We filtered out any missing values for latitude and longitude from all species occurrences. We then cross-checked and cleaned all species occurrences using the function “clean_coordinates” from the package *CoordinateCleaner* (Zizka et al., 2019). 3,621,382 squamate species occurrences and 7,481,576 mammalian species occurrences were retained from the original requested number of occurrences from GBIF.

Using the occurrences obtained above, climatic variables from the online database, WorldClim version 2 (http://www.worldclim.org), annual mean temperature (BIO1) and annual precipitation (BIO12) values were extracted at a ten arc-minutes resolution for each species occurrence using the function “extract” from the package, *raster* (Hijmans 2021). The median mean annual temperature and annual precipitation values were summarized for each species as predictors in the phylogenetic regression. These broad-scale macroecological predictors obtained from range-wide occurrence data are intended to capture range-wide summaries of environmental conditions experienced by a species, as opposed to the microclimatic predictors associated directly with site-of-measurement body temperature data that may be more directly associated with within-individual and within-species variation in thermoregulation. For birds, species’ environmental temperature data were obtained from Cooney et al (2016), which subsetted data to the species breeding range alone. We note that, while more limited, we chose to use validated subsets from Cooney et al (2016) given the inconsistent geographic ranges and challenge of summarizing realized environmental conditions of migratory birds.

### Phylogenetic trees

Phylogenetic trees were taken from published supertrees of major vertebrate groups. The squamate phylogeny was taken from Tonini et al. (2016) from the consensus backbone. The bird phylogeny was gathered from birdtree.org (Jetz et al., 2012). The mammal tree was taken from the synthetic supertree from (Uyeda et al., 2017), which used the Open Tree of life to synthesize a full phylogeny of mammals (Bininda-Emonds et al., 2007; Hinchliff et al., 2015; Meredith et al., 2011) and dated using congruification (Eastman et al., 2013, details provided in Uyeda et al., 2017). The uncertainty introduced by alternative phylogenies was explored by using a hundred posterior sampled trees from the Bayesian analyses of phylogenies of Upham et al. (2019) for mammals, Tonini et al. (2016) for squamates, and Jetz et al. (2012) for birds (Figure S2). Compared to our primary analysis that favored using consensus phylogenies, these alternative topologies are expected to bias phylogenetic regression in opposing directions. Specifically, consensus and taxonomy-supplemented supertrees are expected to contain many polytomies and unresolved tip species and impose lower levels of covariance between tips than the true species tree. Conversely, Bayesian posterior samples of resolved phylogenies may predict erroneously high covariance due to short terminal branch lengths and higher rates of topological error near the tips. Therefore, we examined whether our primary inferences are robust to these two extremes of bias rather than taking Bayesian posterior uncertainty as the sole source of phylogenetic uncertainty.

### Data analysis

We merged trait data with phylogenies for every clade using the function *make.treedata* from the package *treeplyr* (Román Palacios et al., 2021). Taxonomic names were standardized for all datasets prior to merging by using the *tnrs_match_names* function (Hinchliff et al., 2015) accessed through the package *rotl* (Michonneau et al., 2016).

We filtered out all species with missing T_b_ and environmental predictors and retained 500 squamates, 813 mammals, and 476 bird species (Fig. 1). Diel activity, however, is highly phylogenetically conserved, making it amenable to phylogenetic imputation. Using the package *phylopars* (Bruggeman et al., 2009), we imputed both missing tip states and ancestral nodes for squamates, birds and mammals. Tips and nodes were set to the character state with the highest marginal likelihood at each node under the maximum likelihood estimate for the best fitting model among those compared (equal rates, symmetric, and all-rates different Mk models; Lewis 2001).

**Figure 1:**
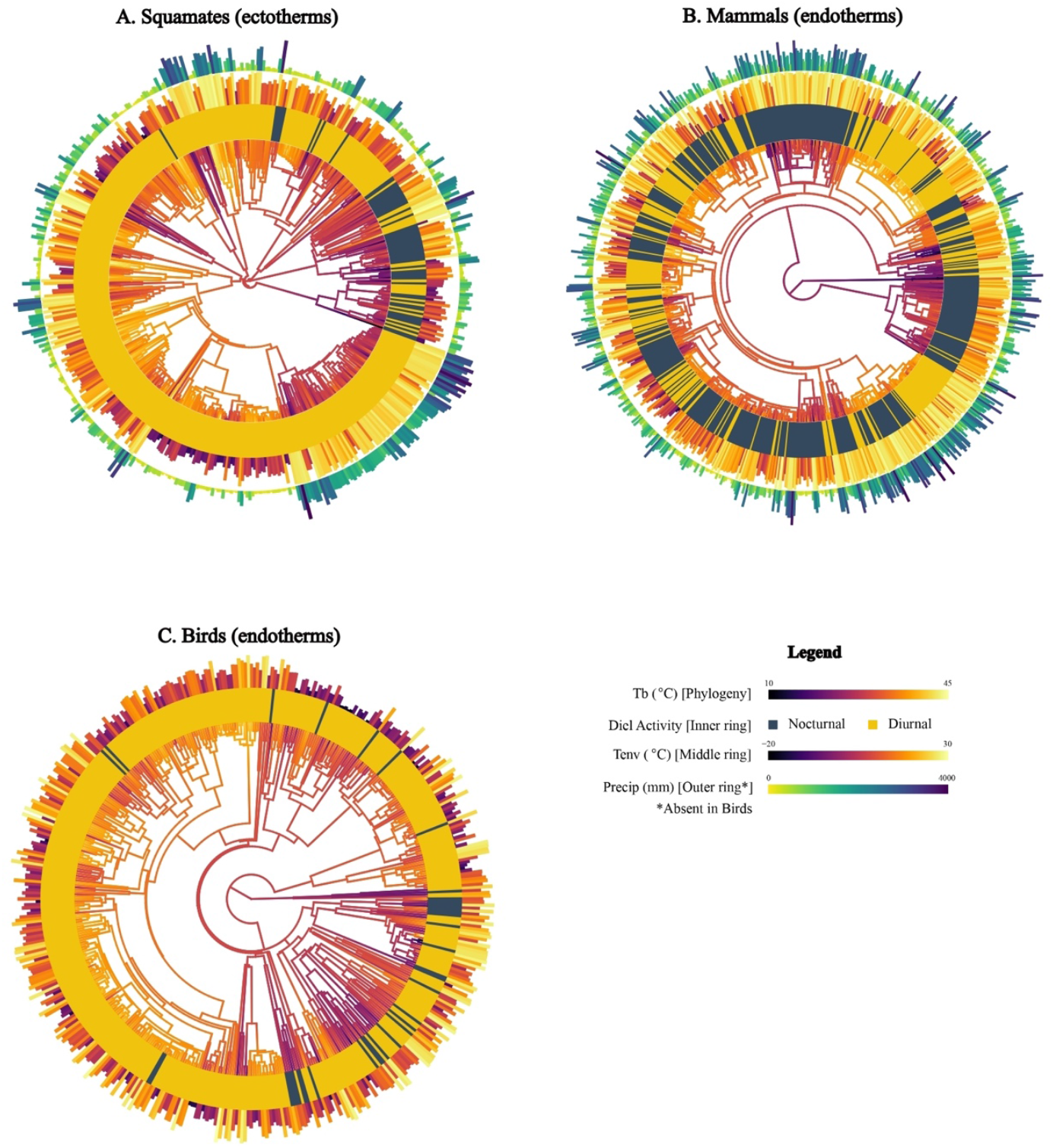
Ancestral state reconstructions of body temperature (for visualization purposes only) for **A**: Squamates, **B**: Mammals, and **C**: Birds indicated by the color of the branches in the middle of each figure. Color grades start at purple for the lowest body temperature to the highest in yellow. The first circle after the phylogeny is the distribution of diurnal (yellow) species and nocturnal (black) species. The second circle represents the environmental temperature; highest temperatures are yellow and with a full bar and the lowest purple. Squamates, **A** and Mammals, **B** have an additional outermost circle for precipitation with the tallest blue bars for the highest precipitation and shorter, pale bars for lower precipitation.

For mammals, we further looked at the effect of rumination and aquatic/terrestrial habitat on body temperature. For the mammal model that included these multiple discrete predictors, we employed the strategy of including them as a single, structured, multistate character (TRD model). Data were coded into three main categories: Terrestrial/Aquatic, Ruminant/non-ruminant, and Nocturnal/Diurnal state. The three states were coded as concurrent (*e.g*., a species could be Terrestrial+Ruminant+Diurnal) and restricted to evolutionary change of one state at a time. We used the matrix function from the package *CorHMM* (Boyko & Beaulieu, 2020) to make the transition rate matrix that fit with the tree, and using the function fit_mk from the package *castor* (Louca & Doebeli, 2018) to estimate the ancestral reconstructions and impute missing data. Matrix set up and assumptions found in the supplementary material Table S2. The resulting discrete predictors were included in *slouch* under stochastic character maps under the reconstructed model.

### Phylogenetic analysis

Analyses were conducted separately for each clade (birds, mammals, squamates) so that we could independently estimate parameters and avoid biases associated with unequal clade sizes. All OU models were implemented by *slouch*, which enables robust estimates of adaptation and inertia as well as the associations between our predictors and response variables. Each independent variable was first tested for its individual influence on body temperature evolution, and then all the variables were analyzed jointly. In all clades, we then selected the focal model for additional analyses based on the model that had the lowest AIC. Further, the *slouch* framework can account for measurement variance and error of species, which otherwise could result in underestimation of values of phylogenetic correlations between species if not accounted for. For species for which we had measurement variance, we provided these. For species with only sample sizes, we pooled the within species standard error from other species. For species data without any estimate of measurement variance, we used the median value from across species for which data were available. The predictor variables; precipitation and environmental temperature were modeled as random covariates evolving by Brownian motion. The model parameters include the half-life, which are valued in millions of years (ma) and the stationary variance; which quantifies the spread of the species’ trait values around the adaptive optima (°C^2^) assuming lineages have reached stationarity around their optimum. In estimation of *slouch*’s OU-BM model, it is typical that half-life and stationary variance covary. To obtain the region of support and quantify uncertainty in our estimates of phylogenetic half-life (*i.e*., inertia) from our best-fitting model, we further used the *slouch* package to generate a likelihood surface around the peak using a grid search to find all regions within 2 log likelihood units of the peak. This surface provides a visualization of the region of support for our estimates of half-life and stationary variance.

## Results

For squamates, we found a positive relationship between body temperature (T_b_) and environmental temperature for both optimal (Slope±SE = 0.358 ± 0.067; R^2^ = 0.13; Fig. 2A; Table 1) and evolutionary regressions (Slope±SE = 0.174 ± 0.032; Fig. 2A; Table 1). The optimal regression for squamates had a steeper and stronger relationship between annual mean temperature and body temperature than the evolutionary regression. We also found a negative relationship between body temperature and precipitation for both optimal (Slope±SE = -0.004 ± 0.001; Fig. 2D; Table 1) and evolutionary (Slope±SE = -2.0×10^−3^ ± 2.0×10^−4^; Fig. 2D; Table 1) regressions within squamates. Consistent with significant phylogenetic inertia (see below), the slopes of the optimal regressions were steeper than the evolutionary regressions. The regime optimum for Nocturnal species was 23.42 ± 1.82 while that of Diurnal species 29.35 ± 1.05.

**Table 1.** Comparison of *slouch* models for the response trait, body temperature (oC), in squamates and mammals (Temperature, Precipitation and Imputed Diel)

|  |  |  |  |  |  | Optimal | Evolutionary |  |
| --- | --- | --- | --- | --- | --- | --- | --- | --- |
| Clade | Predictors<sup>†</sup> | $t_{1/2}^{\ddagger}$ | $\alpha^*$ | $v_y^{\wedge}$ | Intercept (°C±SE) | Slope (±SE)<sup>§</sup> | Slope<sup>§</sup> | $R^2$ |
| Squamates | ***T<sub>env</sub>*+*precip*+*diel*** | 76.71 | 0.01 | 15.57 | Noc: 23.41 ± 1.82<br>Diu: 29.34 ± 1.05 | *T<sub>env</sub>*: 0.36±0.07<br>*precip*: -0.004± 0.001 | *T<sub>env</sub>*: 0.17± 0.03<br>*precip*: -0.0020± 0.0002 | 0.13 |
| Mammals | ***T<sub>env</sub>*+*precip*+*diel*** | 63.30 | 0.01 | 3.55 | Noc: 35.01 ± 0.57<br>Diu: 36.13 ± 0.79 | *T<sub>env</sub>*: -0.024±0.009<br>*precip*: -0.00001± .00009 | *T<sub>env</sub>*: -0.015±0.006<br>*precip*: -0.000007±0.00006 | 0.03 |
| Birds | ***T<sub>env</sub>*+*diel*** | 49.26 | 0.01 | 1.65 | Noc: 38.84 ± 0.69<br>Diu: 40.34 ± 0.33 | *T<sub>env</sub>*: 0.032± 0.048 | *T<sub>env</sub>*: 0.0036 ± 0.0055 | 0.01 |
| Mammals (TRD) | ***T<sub>env</sub>*+*precip*+*TRD*** | 62.56 | 0.01 | 3.50 | TRD: 39.74 ± 3.31<br>TRN: 37.32 ± 4.28<br>TNN: 35.09 ± 0.56<br>TND: 36.15 ± 0.81<br>Aq: 34.27 ± 2.06 | *T<sub>env</sub>*: -0.02±0.01<br>*precip*: -0.0000080±0.000094 | *T<sub>env</sub>*: -0.02±0.01<br>*precip*: -0.000005± 0.000058 | 0.02 |

<sup>†</sup>The values in bold were the chosen models based on AICc values. Predictor codes: *T<sub>env</sub>* = annual mean environmental temperature; *precip* = annual precipitation; Diel = nocturnal/diurnal; TRD = Terrestrial/Aquatic, Ruminant/Non-ruminant, and Diel state (see Methods & Table S2 for state coding and transitions).
<sup>‡</sup>Phylogenetic half-life ( $t_{1/2}$ ) values are in millions of years (my).
\* Rate of adaptation ( $\alpha$ ) values are in $\text{my}^{-1}$ ,
<sup>^</sup>Stationary variance ( $v_y$ ) values are in $^{\circ}\text{C}^2$
<sup>§</sup>The slope (±SE) and intercept (±SE) of the optimal and evolutionary regressions were corrected for measurement error in the body temperature data collected. Slope units for *T<sub>env</sub>* in $^{\circ}\text{C } T_b$ per $1^{\circ}\text{C } T_{env}$ . Units for *precip* in $^{\circ}\text{C } T_b$ per mm precipitation.

**Figure 2.**
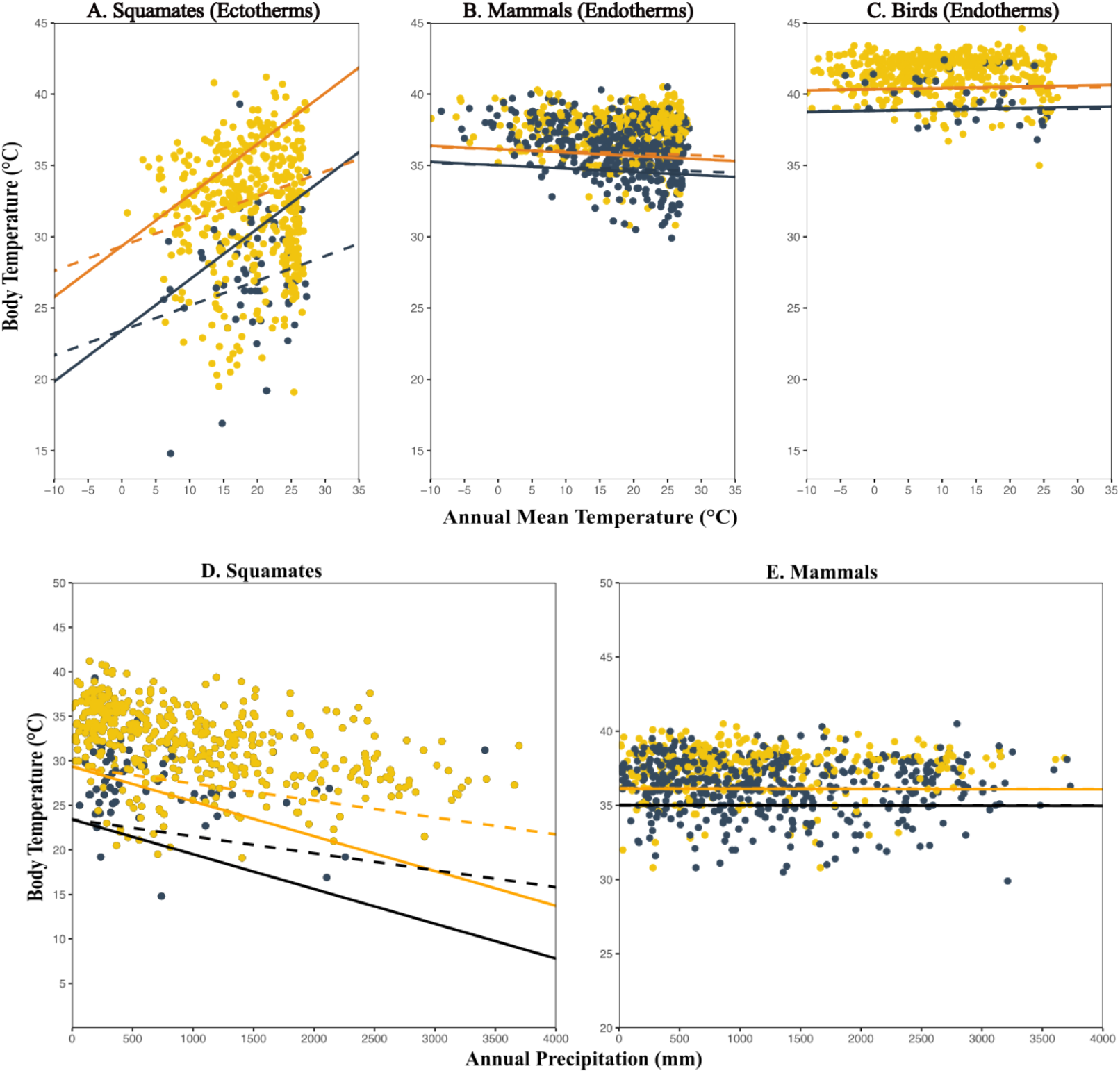
Regressions of body temperature against annual mean temperature for Squamates (**A**), Mammals (**B**), and Birds (**C**), and body temperature against annual precipitation for Squamates (**D**) and Mammals (**E**). Yellow and black points represent diurnal and nocturnal species, respectively, with dashed lines indicating the evolutionary regression. Solid lines indicate the optimal regression, *i.e*., the ideal values of T_b_ if species are allowed enough time to fully adapt to their environments. Note that in endotherms, figures B, C and E, the slopes for the evolutionary and optimal regressions appear similar due to their independence from the environment, despite high estimated levels of inertia.

In contrast, mammals exhibited a statistically significant, yet weak, negative relationship between annual mean temperature and body temperature for both optimal (Slope±SE = −2.366×10^−2^ ± 9.3×10^−3^; R^2^ = 0.0266; Fig. 2B; Table 1) and evolutionary (Slope±SE = -1.46×10^−2^ ± 5.7×10^−3^ Fig. 2B; Table 1) regressions. The optimal regression for annual mean and mammalian body temperatures was slightly steeper than the evolutionary regression. Mammals also showed little to no relationship between annual precipitation and body temperature for both optimal (Slope±SE = -1.156×10^−5^ ± 9.4×10^−5^; Fig. 2E; Table 1) and evolutionary regressions (Slope±SE = -7.13×10^−6^ ± 5.8×10^−5^; Fig. 2E; Table 1). The Regime optima for Nocturnal mammals was 35.01 ± 0.57 and that of Diurnal was 36.13 ± 0.79. Rumination state and aquatic habit to diel state had little effect on the relationship, and the relationship between annual mean temperature and body temperature for both optimal (Slope±SE = -0.02 ± 0.01; Table 1) and evolutionary regressions were indistinguishable (Slope±SE = -0.02 ± 0.01; Table 1). This was also similar to the relationship between annual precipitation and body temperature on both optimal (Slope ≅0.0; Table 1) and evolutionary regression (Slope ≅0.0; Fig. 2D; Table 1). We further subsetted the mammals to only include ruminants (89 species) and again observed little to no relationship between annual mean temperature and body temperature for both optimal (Slope±SE = -5.16×10^−2^ ± 4.27×10^−2^; Table 1) and evolutionary regressions (Slope±SE=-1.07×10^−2^ ± 8.9×10^−3^; Table 1). This was also similar to the relationship between annual precipitation and body temperature on both optimal (Slope±SE = -1.96×10^−4^ ± 4.9×10^−4^; Table 1) and evolutionary regression (Slope±SE = -4.06×10^−5^ ± 1.02×10^−4^; Table 1).

Birds showed a weak, non-significant positive relationship between annual mean temperature and body temperature for both optimal (Slope±SE = 3.16×10^−2^ ± 4.78×10^−2^; Fig. 2C; Table 1) and evolutionary regressions (Slope ± SE = 3.6×10^−3^ ± 5.5×10^−3^; Fig. 2C; Table 1). Birds’ optimal regime was 38.8 ± 0.69 for Nocturnal species and 40.34 ± 0.33 for Diurnal species. The rest of the models for all species can be found in the supplementary materials Table 1.

Despite strong differences in their dependency on environmental temperature and precipitation, all three clades had high levels of phylogenetic inertia as measured by phylogenetic half-life (Squamates: 76.71 my, Mammals: 63.18 my and Birds: 49.26 my). Examination of the likelihood surfaces reveals that all three clades had half-lives that are well above 0, with the lower bound of the support regions beginning after around 40-50 my (Figure 3) and upper bounds extend above hundred-million year timescales (Figure 3). Re-estimation of these values under the posterior distribution of trees for all clades resulted in similar half-life estimates (Supplementary Fig S2).

**Figure 3.**
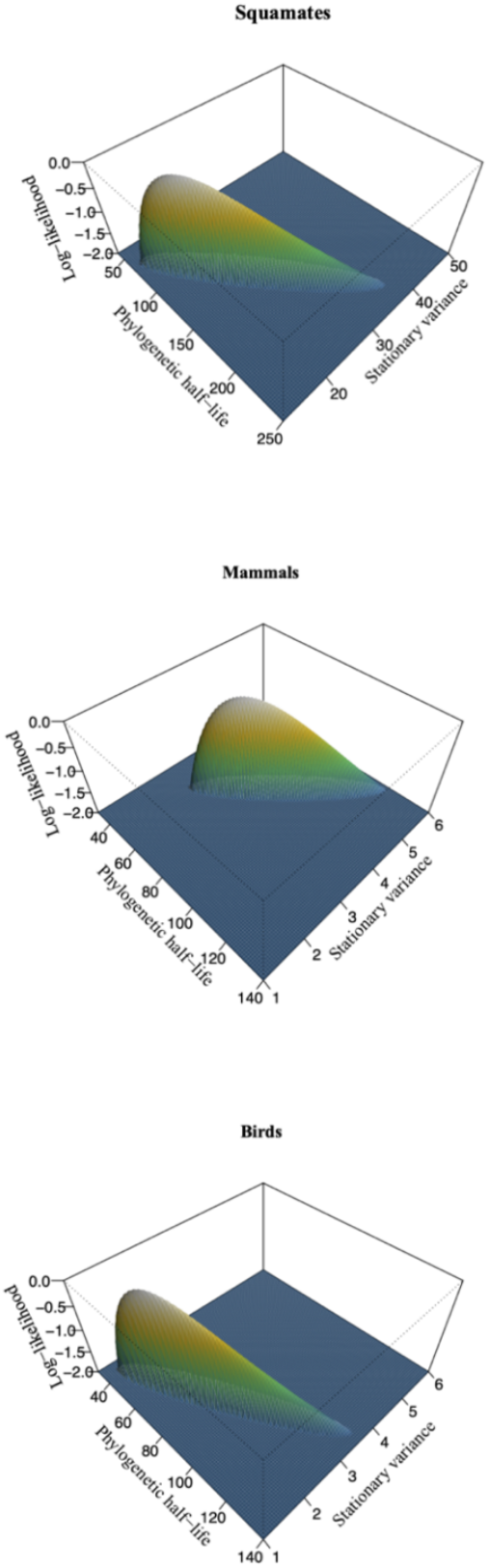
The likelihood surface plots of phylogenetic half-life and stationary variance for Squamates, Mammals and Birds. The elevated areas show the 2-unit support areas for the best model parameters.

## DISCUSSION

Our analyses show that environmental factors, such as annual temperature, precipitation, and diel states (i.e., diurnal and nocturnal), have significant effects on body temperature in squamates. Mammals and birds have decoupled this dependence by regulating their body temperature irrespective of their environments, as evidenced by the weak evolutionary and optimal regressions with environmental variables. Despite differences in thermoregulatory strategies and physiology, body temperatures in endotherms and ectotherms share fundamental similarities in their evolution. Specifically, we observed similarly high levels of phylogenetic inertia, the relationship between the adaptive and optimal values, suggesting similar fundamental constraints on thermal evolvability over macroevolutionary timescales may be shared among clades. Additionally, we found that aquatic/terrestrial habitats and rumination digestive adaptations do not noticeably influence the evolution of body temperature at macroevolutionary scales.

As with previous studies (*e.g*., Clusella-Trullas et al., 2011), we find that environmental temperature, precipitation, and diel state predict thermal physiological parameters in ectothermic tetrapods. Studies using PGLS typically estimated similar relationships between environmental temperature and body temperature in squamates to our estimates. For example, Qu & Wiens (2020) found a 0.12℃ change of body temperature for every 1℃ change in environmental temperature. These findings aligned with our evolutionary regression estimates, which suggest a change of 0.17 ℃ T_b_ with every 1℃ change in environmental temperature. In contrast, we estimate an over two-fold higher relationship in the optimal regression from the adaptation-inertia model. That is, if each lineage is given infinite time to adapt to the environment, we would observe a 0.36 ℃ change in T_b_ for every 1℃ change in environmental temperature. This significantly higher estimate suggests much stronger relationships between environment and body temperature limited by evolutionary constraints and inertia.

While studies on the rates of the evolution of body temperature showed faster rates in ectotherms than endotherms (Moreira et al., 2021), our findings suggest similar levels of trait conservatism— as defined by phylogenetic inertia. When Rolland et al., (2018) modeled niche evolution of endotherms and ectotherms based on their current geographical occurrence, they found endotherms to have the fastest niche evolutionary rates compared to ectotherms. Our results cast new light on this finding, suggesting that higher rates of niche evolution are achieved by decoupling the dependence to the environment of endotherms’ physiology. In this context, it is notable that endothermy can be interpreted as a physiological mechanism to escape the organismal dependency on the environment for setting the thermal optimum of body temperature, while simultaneously increasing their diversity of environmental thermal niche space.

When organisms face environmental changes, they often respond by evolving behavioral and/or physiological mechanisms to maintain a preferred body temperature. Endotherms typically rely on physiological thermoregulation whereas ectotherms primarily use behavioral strategies to cope with changing environments. This reliance on behavior, possibly due to the relative ease of evolving behavioral traits (Blomberg et al., 2003), can buffer organisms from selective pressures. As a result, directional selection is weakened, and evolutionary change is slowed. Essentially, behavioral thermoregulation can lead to retaining of ancestral traits on current niches, creating a lag in adaptation. This phenomenon is known as the Bogert effect (Huey et al. 2003; Muñoz 2022) and explains why we don’t observe an instant adaptation to the current environment: species track their ancestral niches and employ behavior to maintain a preferred body temperature (Anderson & Wiens, 2017).

While endothermic organisms, by definition, require less behavioral thermoregulation to maintain optimal body temperatures, an expanded view of endothermy itself suggests it can be viewed as a “Bogert effect-like” adaptation that allows organisms to escape from the fundamental constraints of thermal evolvability that appear shared among lineages. Specifically, we can view endotherms as decoupling their dependence on the environment and deploying physiological adaptations to maintain their body temperature within a narrow range to a preferred body temperature against a broader range of environmental temperatures. This hypothesis is supported by the observation that T_b_ is a trait that is still adapting to its optimal temperature, with high half-lives just like ectotherms, but with dramatically altered optimal regression slopes (Table 1). On the other hand, the common pattern of macroevolutionary stasis observed in both endotherms and thermoregulating ectotherms might reflect not only shared decoupling of body temperature evolution and environmental temperature, but rather shared evolutionary constraints. In both groups, individuals are composed of thousands of proteins that must undergo coordinated evolution—a “correlated progression” (Kemp 2006, 2007; Labra et al. 2009)—to respond to thermal selection pressure, as changes in organismal body temperature will have cascading effects on all expressed proteins, which undoubtedly vary in their thermal performance profiles (Rezende & Bozinovich 2019, Rahban et al. 2022). If this is true, even ectotherms with less pronounced thermoregulatory behavior should experience evolutionary constraints. Mechanistic, microevolutionary models of body temperature evolution will clarify the expected consequences of the correlated progression of proteins. The results of our study suggest shared macroevolutionary constraints on thermal biology may exist in both endotherms and ectotherms, implicating shared biochemical and genetic constraints underlying these limits on macroevolutionary change.

The stronger physiological yoke to the environment observed in ectotherms than in endotherms may help explain the larger ranges and higher diversity of habitats occupied by the latter (Rolland et al. 2018). Further, the strong relationship we found between body temperature and diel states, especially in ectotherms where we found a ∼5 ℃ difference in optimal states between Nocturnal and Diurnal states and a ∼1 ℃ difference in endothermic species, is consistent with the findings of Moreira et al. (2021). Ectothermic species rely on the environment to control their body temperature and are active when environmental temperatures are close to their preferred range. The stasis in physiological traits such as body temperature is best defined as a restricted adaptation of a species to their current environment or, more generally, the similarities in body temperature values in species that otherwise occur at a broad range of environmental variation, reflecting a species’ resistance to depart from their ancestral state (Eldredge, 2024; Voje, 2016; Williams, 1992).

We could imagine a hypothetical organism capable of achieving maximal fitness would do so by adapting to their environment instantly. However, biological constraints can result from macroevolutionary associations between external drivers of selection, such as environmental temperature, and complex architecture of organismal traits such as body temperature (Hansen et al., 2008). Specifically, such traits are not expected to instantaneously fit to the evolutionarily optimal relationship. While both endotherms and ectotherms are expected to evolve to adapt to their environment to minimize the costs of behavioral and physiological thermal regulation, the signal to adapt comes from the external environment, which continuously changes stochastically and can generate lags such that organismal traits trail behind the optimum set by the environment (Hansen et al., 2008, Tattersall et al., 2012). By modeling change in the ancestral states of these environmental predictors from their current values, we estimated that body temperature does indeed evolve with significant adaptive lag (*i.e*., phylogenetic inertia). Labra et al. (2009) estimated low inertia in *Liolaemus* lizards, albeit with much smaller sample sizes and wide support regions that span our estimates. By analyzing multiple selective factors in a framework that accounts for this inertia, we obtain estimates of dependency between environmental variables and body temperature that are much stronger than previous estimates (Medina et al., 2012; Qu & Wiens, 2020), which assumes instantaneous adaptation with phylogenetic covarying residuals. Our approach simultaneously reveals strikingly similar levels of inertia across both endotherms and ectotherms, suggesting common constraints on thermal adaptation.

## Concluding remarks

Our findings unify observations of body temperature evolution and find similar levels of inertia of body temperature evolution in both endotherms and ectotherms. We propose that this represents a universal constraint on thermal adaptation in vertebrates. As with physiological studies that observe the preference for behavioral adaptation to explain the relatively static physiological traits, (*i.e*., the Bogert effect), we extend these behavioral adaptations to the concept of endothermy itself. In other words, endotherms can be seen as an adaptation maintaining physiological traits with low-evolvability at their functional optima, unlinking them to variable and changing selective environments and allowing them to track faster moving climatic optima, explaining faster rates of niche evolution in endotherms. Conversely, our macroevolutionary data suggest particularly strong limits to how quickly ectotherms can achieve optimal body temperatures in rapidly changing thermal environments.

