## Supplementary material for "Universal phylogenetic inertia in body temperature evolution across endothermic and ectothermic tetrapods": Supplemental Document.docx

**Table S1.** All models explored with all the predictor variables. Best models are in bold. Note that all the best-fitting models are comprised of both environmental variables of temperature and precipitation.

|  |  |  |  |  | **Optimal** | | **Evolutionary** |  |  |
| --- | --- | --- | --- | --- | --- | --- | --- | --- | --- |
| **Species Group** | **Predictor(s)** | **t1/2**  **(mya)** | **α**  **(mya-1)** | **νy (°C²)** | **Intercept (±SE)** | **Slope (±SE)** | **Slope** | ***R*²** | **AICc** |
| Squamates | Temp | 42.14 | 0.02 | 20.78 | 29.01± 0.88 | 0.12± 0.05 | 0.08± 0.04 | 0.01 | 2,701.64 |
| Squamates | Precip | 47.37 | 0.01 | 19.21 | 32.17± 0.68 | -0.003± 0.00 | -0.002± 0.00 | 0.09 | 2,656.30 |
| Squamates | Temp + Precip | 99.04 | 0.01 | 19.03 | 28.33 ± 1.11 | precipmm -0.004± 0.00  tempC 0.41± 0.079 | precipmm 0.002 ± 0.00  tempC 0.17 ± 0.03 | 0.11 | 2,554.23 |
| **Squamates 500** | **Temp + Precip+Diel** | **76.71** | **0.01** | **15.57** | **0 23.42 ± 1.82 1 29.35 ± 1.05** | **precipmm -0.004± 0.00**  **tempC 0.36± 0.07** | **precipmm -0.002 ± 0.00 tempC 0.17± 0.03** | **0.13** | **2,546.70** |
| Mammals | Temp | 31.71 | 0.02 | 3.46 | 35.73 ± 0.3 | -0.02± 0.01 | 0.01± 0.01 | 0.01 | 2,675.51 |
| Mammals | Precip | 32.72 | 0.02 | 3.52 | 35.64 | -0.0002±0.00 | 0.00±0.00 | 0.01 | 2,673.50 |
| Mammals | Temp + Precip | 67.56 | 0.01 | 3.74 | 35.15 | Precipmm 0.00±0.00 tempC  0.02±  0.01 | precipmm 0.00 ±0.00 tempC 0.01±0.01 | 0.01 | 2,558.09 |
| **Mammals 813** | **Temp + Precip + Diel** | **63.18** | **0.01** | **3.54** | **0 35.00 ±0.57 1 36.18 ± 0.79** | **precipmm 0.00± 0.00 tempC 0.02 ± 0.01** | **precipmm 0.00 ± 0.00** **tempC 0.01 ± 0.01** | **0.01** | **2,557.22** |
| Birds | Temp | 56.62 | 0.01 | 1.84 | 40.18 ± 0.37 | 0.01± 0.01 | 0.004±0.005 | 0 | 1,330.00 |
| **Birds 476** | **Temp + Diel** | **49.26** | **0.01** | **1.65** | **0 38.84± 0.69**  **1 40.34 ± 0.33** | **0.01± 0.01** | **0.004**±**0.005** | **0.01** | **1,330.00** |
| Ruminants 89 | Temp + Precip | 60.61783 | 0.01143471 | 0.9056694 | 38.63 ± 0.46 | 0.00±0.004  0.05± 0.043 | 0.00 ± 0.00  0.0 ± 0.01 | 0.0266 | 185 |
| TRD 813 | Temp + Precip | 62.555 | 0.0110806 | 3.4989 | 1 39.74 ± 3.3  2 37.32 ± 4.3  3 35.09± 0.6  4 36.15± 0.8  5 34.27± 2.1 | Precip 0.00± 0.00  Temp 0.02 ± 0.01 | precipmm 0.00 ± 0.00  tempC 0.02 ± 0.01 | 0.02 | 2,561.00 |

**^†^**The values in bold were the chosen models based on AICc values.

**^‡^**Phylogenetic half-life (t_1/2_) values are in millions of years (mya).

^*^ Rate of adaptation (α) values are in mya^-1^,

^^^Stationary variance (ν_y_) values are in ^o^C^2^

^§^The slope (±SE) and intercept (±SE) of the optimal and evolutionary regressions were corrected for measurement error in the body temperature data collected.

**Table S2.** TRD model. Terrestrial/Aquatic, Ruminant/Non-Ruminant and Diel/Nocturnal transition matrix. The three states were coded as concurrent states (eg. a species could be Terrestrial+Ruminant+Diurnal) with unequal transition rates to another state and restricted to evolutionary change of one state at a time e.g a Terrestrial+Ruminant+Diurnal (TRD) species cannot transition to Terrestrial+Non-Ruminant+Nocturnal (TNN) species in one step.

|  | TRD | TRN | TNN | TND | Aq |
| --- | --- | --- | --- | --- | --- |
| TRD | - | ⍺Noc | 0 | 0 | ⍺A |
| TRN | ⍺Di | - | 0 | 0 | ⍺A |
| TNN | 0 | ⍺Ru | - | ⍺Di | ⍺A |
| TND | ⍺Ru | 0 | ⍺Noc | - | ⍺A |
| Aq | 0 | 0 | 0 | 0 | - |

Further model assumptions include:

a. Diel activity and rumination were not relevant to aquatic species; therefore, the state collapsed to just Aquatic (Aq) irrespective of other states for all transitions from Terrestrial.
b. Organisms cannot transition back to a terrestrial state from an aquatic state.
c. Rumination state is irreversible

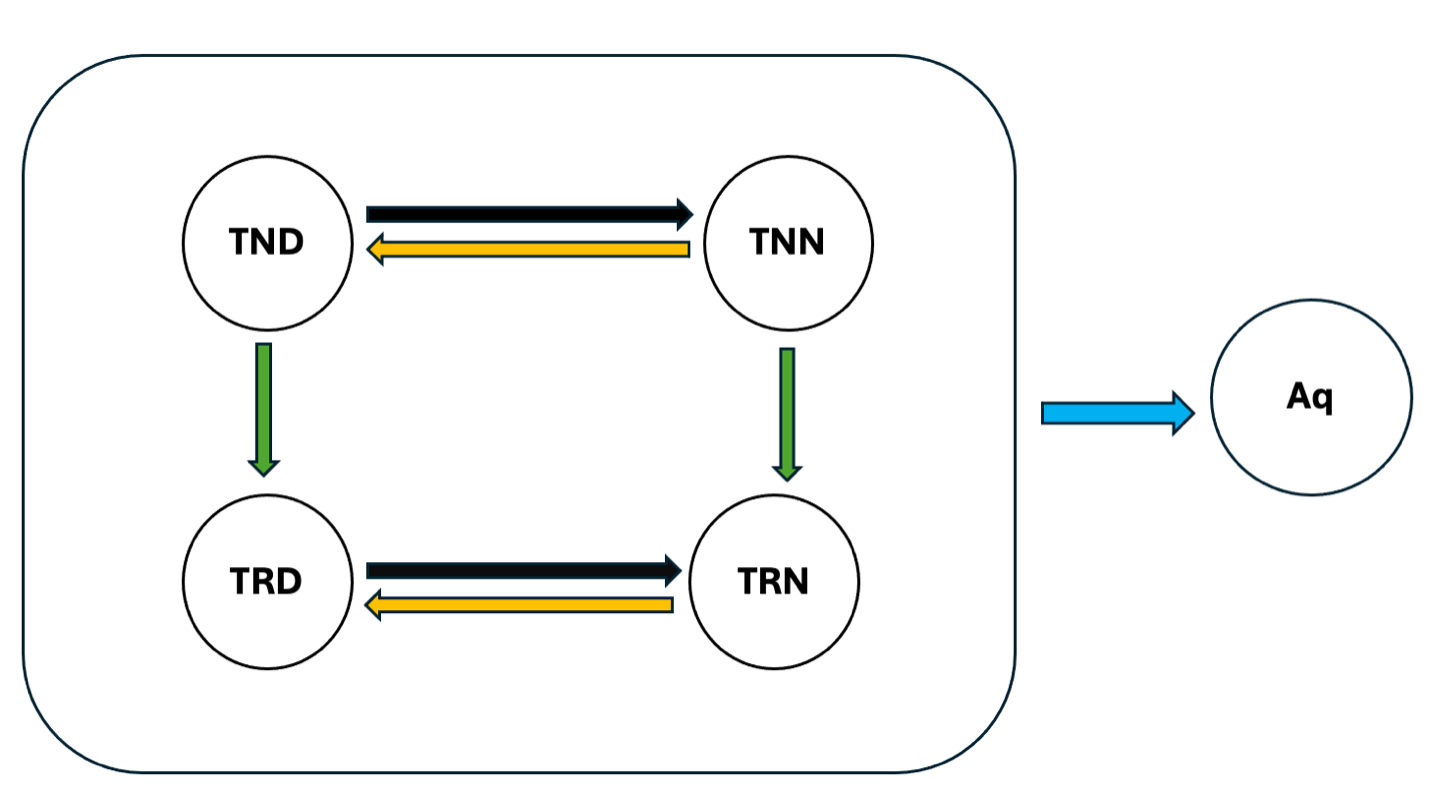
 **Figure S1:** Figure to alternatively visualize the fitted transition matrix in Table S2. Different arrow colors for different rates from one state to another. Black are changes to Nocturnal, Yellow to Diurnal, Green to Ruminants, and Blue to an Aquatic state (e.g TND represents Terrestrial-Non Ruminant-Diurnal mammal). Each of the states in the box can change to aquatic at a shared transition rate. We disallow back-transitions from ruminants and aquatic states since these transitions are not expected to exist in the mammal phylogeny, and thus avoids having transition parameters linked to unobserved transitions.

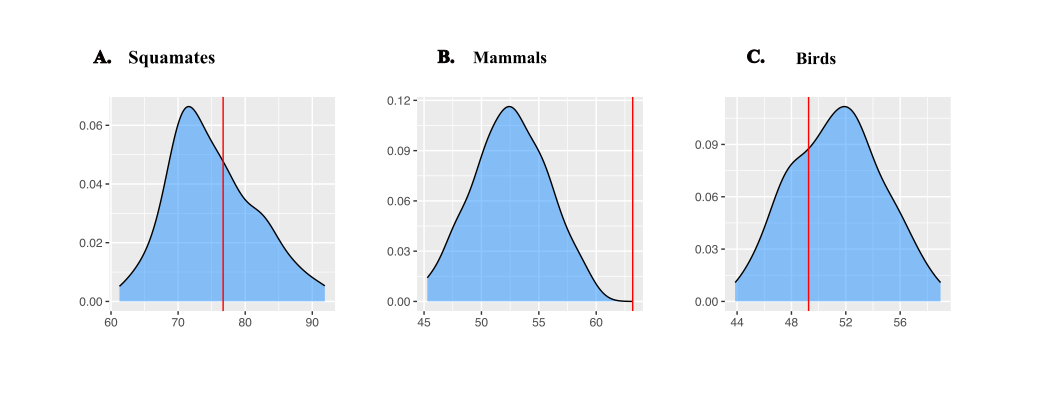
**Figure S2**: Figure showing the distributions of maximum likelihood values of half-lives estimated from a hundred alternative phylogenies for (A) Squamates, (B) Mammals and (C) Birds. The primary estimates presented in the main text are given by the vertical red lines. Note that the tree we used for our analysis in mammals has younger nodes than the distribution of the alternative phylogenies, hence the bias of the model to a higher half-life. For mammals, we found that half-lives estimated from the (Upham et al., 2019) phylogeny had overall shorter half-lives vs. our original analysis due to using a different phylogeny than (Uyeda et al., 2017). This occurs because the Upham phylogeny has relatively younger node ages than the Uyeda et al. 2017 tree (median node age in Upham et al.: 7.8 my, root age: 181.4my vs. median node age in Uyeda et al. 2017: 11.8my, root age: 215.6my) for the same overlapping set of taxa. In addition, Upham et al. contained relatively more of the youngest node splits below a million years (3.9% of nodes vs. 2.8% of nodes), potentially contributing to biological sources of “tip fog” (Beaulieu & O’Meara, 2024), which further decreases apparent phylogenetic signal and increases estimates of half-life. Nevertheless, note that the distributions shown here represent point estimates for each tree while estimation uncertainty given the likelihood surfaces (e.g. Figure 3) is much greater, with longer tails toward longer half-lives.
